# Genetic editing of *CISH* enhances T cell effector programs independently of immune checkpoint cell surface ligand expression

**DOI:** 10.1101/2021.08.17.456714

**Authors:** Elisa Arthofer, Krishnendu Chakraborty, Lydia Viney, Matthew J Johnson, Beau R. Webber, Branden S. Moriarity, Emil Lou, Modassir Choudhry, Christopher A. Klebanoff, Tom Henley

## Abstract

PD-1 acts as a negative regulator of T cell-mediated immune responses in the setting of persistent antigen expression, including cancer and chronic pathogen infections. Antibody-mediated blockade of the PD-1/PD-L1 axis benefits a subset of patients with highly immunogenic malignancies; however, many patients fail to respond due to a requirement for expression of the cell surface ligand PD-L1 within the tumor microenvironment. CISH is a member of a new class of intra-cellular immune checkpoint molecules that function downstream of the T cell receptor to regulate antigen-specific effector functions, including reactivity to cancer neoantigens. Herein, we employed multiplex CRISPR editing of primary human T cells to systematically compare the function of *CISH* deletion relative to *PDCD1* (the gene encoding PD-1) and/or *VSIG9* (the gene encoding TIGIT) in a model of neoantigen-mediated cancer cell cytolysis. PD-1 and TIGIT disruption enhanced cytolytic activity exclusively in the setting of high PD-L1 expression. In contrast, CISH inactivation enhanced antigen-specific cytolysis of tumor cells regardless of PD-L1 expression, including outperforming PD-1 and TIGIT disruption even in the presence of high PD-L1 tumor cells. Furthermore, we observed a synergistic increase in tumor cell killing when CISH and PD-1 or TIGIT are inactivated in combination, supporting the notion that these immune checkpoints regulate non-redundant pathways of T cell activation. Together, these data demonstrate that the intra-cellular immune checkpoint protein CISH can potentially enhance anti-tumor responses against a broad range of cancer types regardless of PD-L1 biomarker status.

## MAIN

T cells play a crucial role in immune-mediated tumor clearance by recognizing tumor cells via their T cell receptors (TCRs) to elicit a program of targeted destruction culminating in cancer cell lysis^1^. A critical requirement for improving clinical responses to immunotherapy lies in enhancing the functional avidity of cancer antigen-specific T cells^2, 3^. Thus, the field of immune checkpoint inhibition has developed to advance the clinical outcomes of anti-cancer therapies via modalities to inhibit T cell immune checkpoints and reduce the immunosuppressive effects these cells encounter^4^. Therapeutic blockade of immune checkpoint receptors or their associated ligands, such as Programmed Cell Death Protein 1 (PD-1) and PD-L1, can trigger the regression of diverse cancer types. These findings underscore the potential of targeted approaches to enhance and prolong antigen-specific T cell responses^5, 6, 7, 8, 9, 10^.

The interaction of PD-1 with its ligands, programmed death 1 ligand (PD-L1) and PD-L2, inhibit T cell activation, suppress proliferation, and limit anti-tumor effector functions through a co-inhibitory signaling pathway^11, 12^. FDA approved monoclonal antibody (mAb) inhibitors of PD-1 and PD-L1 have shown impressive clinical outcomes in a subset of patients resulting in durable tumor regression and extended progression free survival^13, 14^. Despite these clinical successes, most patients show no response to blockade of the PD-1/PD-L1 axis. Several mechanisms have been identified that underly this resistance, most notably a requirement for cells in the tumor microenvironment (TME) to express PD-L1^5, 15, 16 17^. PD-L1 expression within the TME can vary between patients and low or absent expression can render the patient unresponsive to anti-PD-1/PD-L1 mAbs^18, 19^. Thus, there remains a critical need to identify and therapeutically modulate immune checkpoint pathways that function independently of specific cell surface ligands.

The cytokine induced SH2 protein CISH is a recently identified cancer immune checkpoint that functions as a negative modulator of TCR signaling and cancer neoantigen recognition. Unlike PD-1, CISH is an intracellular protein that negatively regulates antigen-specific cytokine release and T cell expansion via its capacity to bind PLC-γ1, a proximal mediator of TCR complex signaling, for targeted proteasomal degradation.^20, 21, 22, 23^. Germline deletion of *Cish* in mouse tumor-specific CD8^+^ T cells promotes their expansion and cytokine polyfunctionality resulting in increased durable regression of established melanoma lesions^22^. Ablation of *CISH* in human TIL cells increases antigen-specific T cell proliferation, TCR functional avidity, and neoantigen reactivity^20^. Additionally, CISH has recently been shown to play an important role in negatively regulating Natural Killer (NK) cell persistence and *in vivo* anti-tumor activity by suppressing activation downstream of the IL-15 receptor^24, 25, 26^.

CISH is distinct from conventional cell surface immune checkpoint receptors. Unlike molecules such as PD-1 and cytotoxic T-lymphocyte-associated protein 4 (CTLA-4) that operate by binding to tumor cells ligands (PD-L1 and CD80 respectively), the CISH-PLC-γ1 interaction occurs within the intra-cellular compartment downstream of the TCR. Although inactivation of cell-surface immune checkpoints may be achieved through mAb-based targeting, intra-cellular signaling molecules such as CISH have historically remained unreachable (and thus undruggable). However, the advent of targeted gene editing tools, such as CRISPR, now permit the precise, irreversible, and efficient inactivation of CISH in human T cells^20^. The potential of this novel immune checkpoint target to improve T cell therapies for solid cancers is now being investigated in patients using CRISPR engineered CISH knockout tumor infiltrating lymphocytes (TIL) (NCT04426669).^27^

In a pre-clinical murine model, we recently demonstrated that CISH knockout results in enhanced tumor regression when combined with PD-1 mAb blockade^20^. In the present manuscript, we now seek to understand the functional relationship between these two immune checkpoint targets further. To test for potential synergy, we developed a highly efficient multiplex CRISPR editing approach for use in primary human T cells. We targeted multiple immune checkpoint genes simultaneously, allowing us to compare the CISH knockout phenotype with that of PD-1 and the evolving cell surface immune checkpoint, T cell Immunoreceptor with Ig and ITIM domains (TIGIT). Analysis of the effector function of immune checkpoint-deficient T cells showed that inactivation of CISH leads to an enhanced program of T cell activation, memory formation, and antigen-specific cancer cell cytolysis that individually is superior to, and in combination synergistic with, PD-1 and TIGIT disruption. Importantly, the benefit of CISH disruption occurred independently of PD-L1 expression, a finding that contrasts with the function of both PD-1 and TIGIT disruption. Together, our findings demonstrate an important role for CISH in controlling T cell responses to cancer in a manner that is independent of PD-L1/PD-L2 ligand expression. These results establish a unique and non-overlapping role for CISH compared to canonical cell surface immune checkpoint targets such as PD-1/PD-L1.

## RESULTS

### Multiplex gene editing enables the evaluation of CISH and other immune checkpoints in regulating antigen-specific T cell function

To evaluate the immune function of CISH in relation to PD-1 and TIGIT in primary human T cells, we developed a CRISPR/Cas9-based strategy for efficient multiplexed gene editing followed by a functional evaluation using a real-time cancer cell cytolytic assay (**Fig. 1a**). We identified multiple guide RNAs targeting the *CISH, PDCD1*, and *VSIG9* (the gene encoding TIGIT) loci that resulted in a significant reduction in expression of the corresponding proteins **(Fig. 1b, c, & d)**. A high level of genetic knockout enabled us to assess the phenotypic and functional consequences of inactivating these immune checkpoints in a head-to-head-to-head fashion. To measure T cell function, we simultaneously introduced a double-strand break within the TCR alpha locus (*TRAC*) combined with adeno-associated virus (AAV) delivery of a DNA repair template to introduce a previously described recombinant TCR specific for the KRAS(G12D) shared cancer neoantigen^28^. Integration of the exogenous KRAS(G12D) TCR occurred at high frequency, as measured by expression of the murine TCRβ constant chain (mTCRβ), while simultaneous CRISPR targeting of the *TRAC* locus removed expression of the endogenous TCR (**Fig. 1e)**. Transgenic TCR integration at the endogenous *TRAC* locus afforded more robust TCR expression than targeting the *AAVS1* safe-harbor site. Stable levels of TCR expression was observed over three-weeks of *ex vivo* T cell culture **(Fig. 1f)**. Overall, this multiplex CRISPR/AAV platform enabled the generation of a pool of edited T cells in which at least 50% have lost expression of two immune checkpoint genes while simultaneously introducing an antigen-specific TCR. This allowed a comparative investigation into the functional impact of immune checkpoint modulation on human T cell biology.

**Figure 1:**
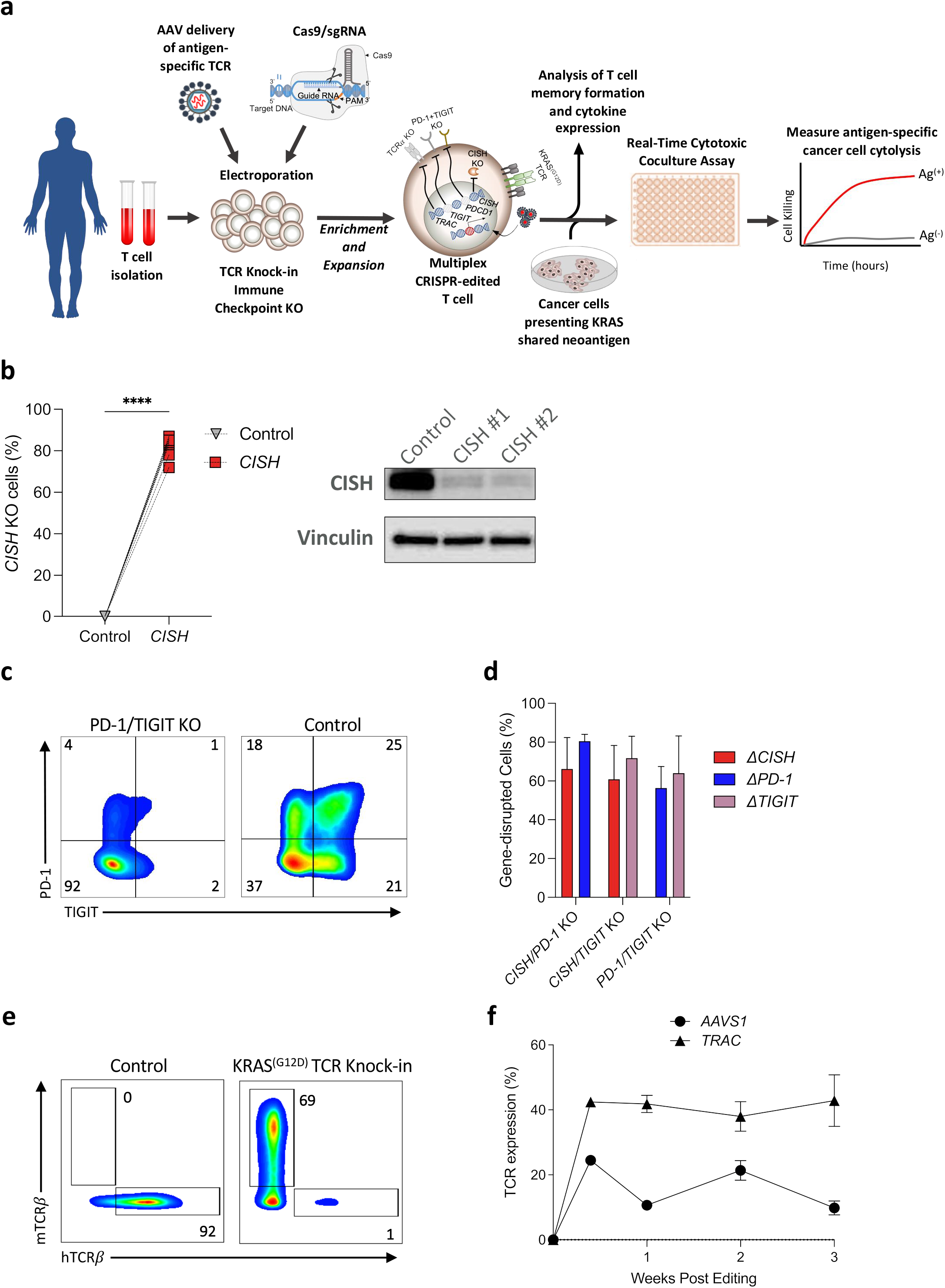
Multiplex CRISPR/rAAV editing of *CISH* and KRAS(G12D) TCR integration in primary human T cells. **(a)** Schematic diagram of a multiplex CRISPR/rAAV genome engineering and cancer cell cytolysis assay platform for primary human CD8^+^ T cells. **(b)** Efficient disruption of the intra-cellular checkpoint gene *CISH* measured on DNA level by Sanger sequencing and reduced CISH protein expression measured by Western blot. **(c)** T cell surface expression of immune checkpoint genes PD-1 and TIGIT measured by flow cytometry with or without multiplex CRISPR editing. **(d)** The frequency of CD8^+^ T cells with disrupted immune checkpoint genes after simultaneous multiplex editing. **(e)** Targeting of the *TRAC* locus for rAAV-mediated insertion of the recombinant KRAS(G12D)-specific TCR results in loss of endogenous TCR expression while enabling high expression of the exogenously introduced TCR. **(f)** Comparison of recombinant TCR expression over 3 weeks following CRISPR/rAAV engineering of primary human CD8^+^ T cells when integrated into the either the *TRAC* or *AAVS1* locus. Statistical significance was determined by either student t test or ANOVA for repeated measures; *P<0.05, **P<0.01, ***P<0.001, ****P<0.0001. All data are representative of at least three independent experiments. Error bars represent mean +/− SEM.

### CISH inactivation enhances T cell activation, cytokine production and the formation of effector memory cells

Given the intra-cellular nature of CISH and its capacity to attenuate proximal TCR signaling, we hypothesized that disruption of CISH in naïve human T (T_N_) cells derived from the peripheral blood would result in an enhanced program of TCR-mediated effector functions. We first analyzed the formation of distinct T cell memory subsets upon anti-CD3/CD28 stimulation of CD8^+^ T_N_ cells by measuring expression of the memory markers CD45RA and CD45RO. We categorized the cell populations as being either T_N_/T stem cell memory (T_SCM_) (CD45RO^-^, CD45RA^+^) or conventional memory phenotype (CD45RO^+^, CD45RA^-^). CISH knockout T cells showed a significant increase in the transition to a conventional memory T cell phenotype when compared to control T cells (**Fig. 2a**). Further delineation of the T cell memory phenotype by analysis of the lymphoid-homing marker CD62L within the CD45RO^+^ population revealed that CISH inactivation elevated the formation of an effector memory T (T_EM_) cell phenotype (CD62L^-^CD45RO^+^CD45RA^-^) (**Fig. 2a & b**).

**Figure 2:**
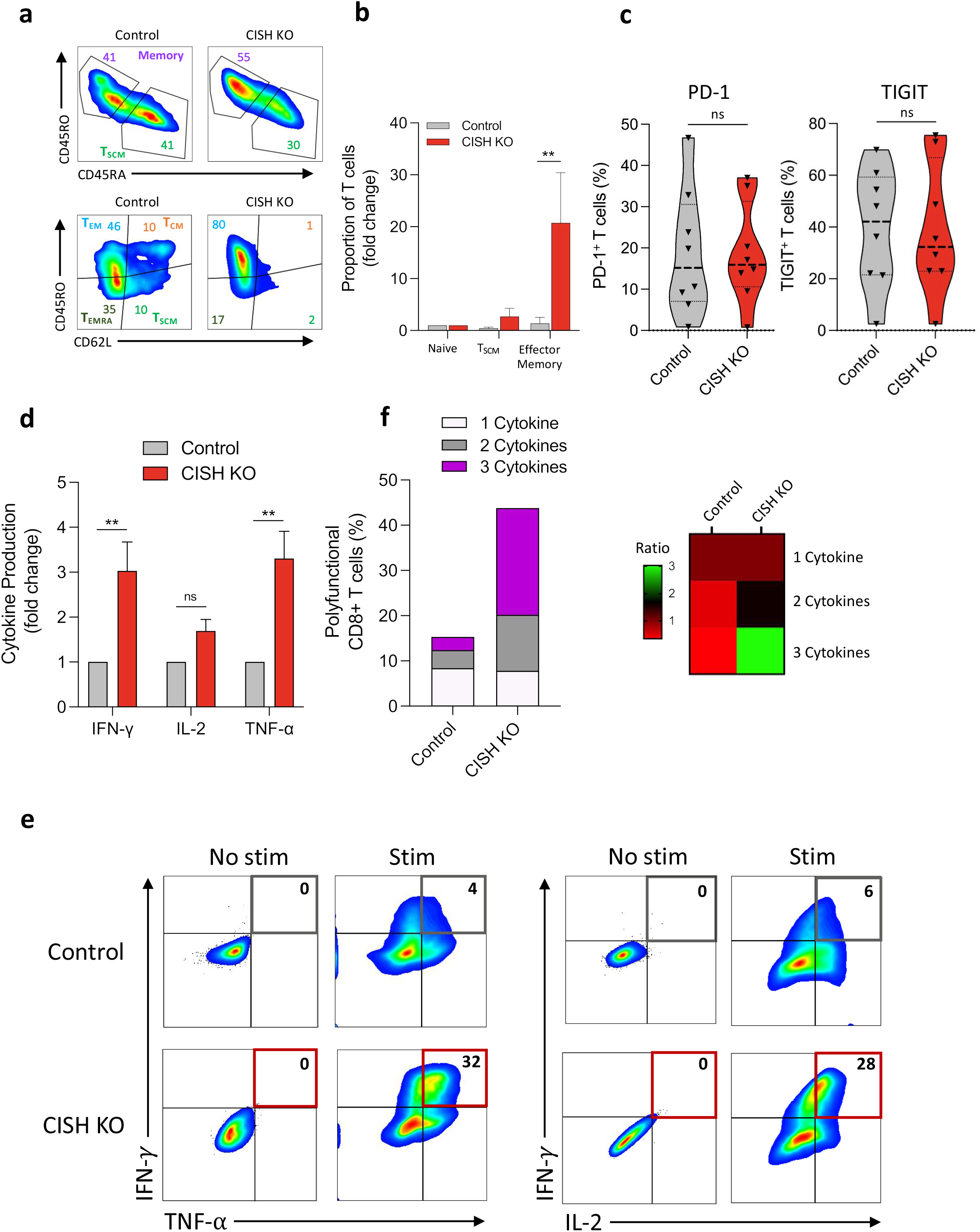
Inactivation of CISH in human T cells enhances T cell effector function upon TCR signaling. **(a)** Knockout (KO) of CISH in CD8^+^ T cells increases the proportion of cells with a memory phenotype upon anti-CD3/CD8 stimulation (upper panels) and the effector memory proportion T_EM_ (lower panels). **(b)** Quantification of changes in memory phenotypes in CD8^+^ T cells in control and CISH-knockout as in (a). **(c)** Expression of inhibitory receptors PD-1 and TIGIT is similar between CISH KO and control T cells. **(d-e)** Knockout of CISH in CD8^+^ T cells significantly increases the magnitude of effector cytokine production and the frequency of T cells expressing 2 or 3 cytokines as measured by intra-cellular cytokine staining (ICS). **(f)** CISH knockout in CD8^+^ T cells increases the total number of polyfunctional CD8^+^ T cells after TCR stimulation via anti-CD3/CD28 beads. In addition, CISH knockout elevates the ratio of T cells expressing 1:2:3 cytokines. For polycytokine visualization one representative donor is shown. Statistical significance was determined by either student t test or ANOVA for repeated measures, *P >0.05, **P>0.01, ***P>0.001, ****P>0.0001. All data are representative of at least three independent experiments. Error bars represent mean +/− SEM.

Expression of the co-inhibitory PD-1 and TIGIT receptors was similar between control and CISH knockout peripheral T cells (**Fig. 2c**). TCR stimulation alone was sufficient to reveal a stronger induction of cytokine secretion by CISH knockout versus control T cells and a significant increase in IFN-γ and TNFα expression (**Fig. 2d**) and increased cytokine polyfunctionality (**Fig. 2e & f**). No cytokine secretion by CISH knockout T cells was observed in the absence of TCR stimulation. Taken together, these data show that TCR stimulation of human CISH knockout T cells results in a significant elevation in the formation of activated memory T cells with increased production of multiple cytokines.

### PD-1 or TIGIT knockout T cells fail to enhance T cell function following TCR stimulation

T cells deficient for either PD-1 or TIGIT showed no increase in the formation of T_EM_ cells in response to anti-CD3/CD28 stimulation, in contrast with CISH KO cells **(Fig. 3a & b)**. Furthermore, lack of PD-1 or TIGIT also had no impact on the production of cytokines or cytokine polyfunctionality, whereas CISH significantly elevated IFN-γ, TNFα, and the proportion of T cells expressing 2 or more cytokines (**Fig. 3c & d)**. As seen with CISH knockout T cells, lack of PD-1 did not increase expression of TIGIT, and conversely lack of TIGIT did not increase expression of PD-1 **(Fig. 3e)**. The lack of an increase in functional response when PD-1 and TIGIT are deleted in T cells suggests that stimulation of the TCR alone is insufficient to enhance T cell activation in the absence of expression for cell surface immune checkpoints.

**Figure 3:**
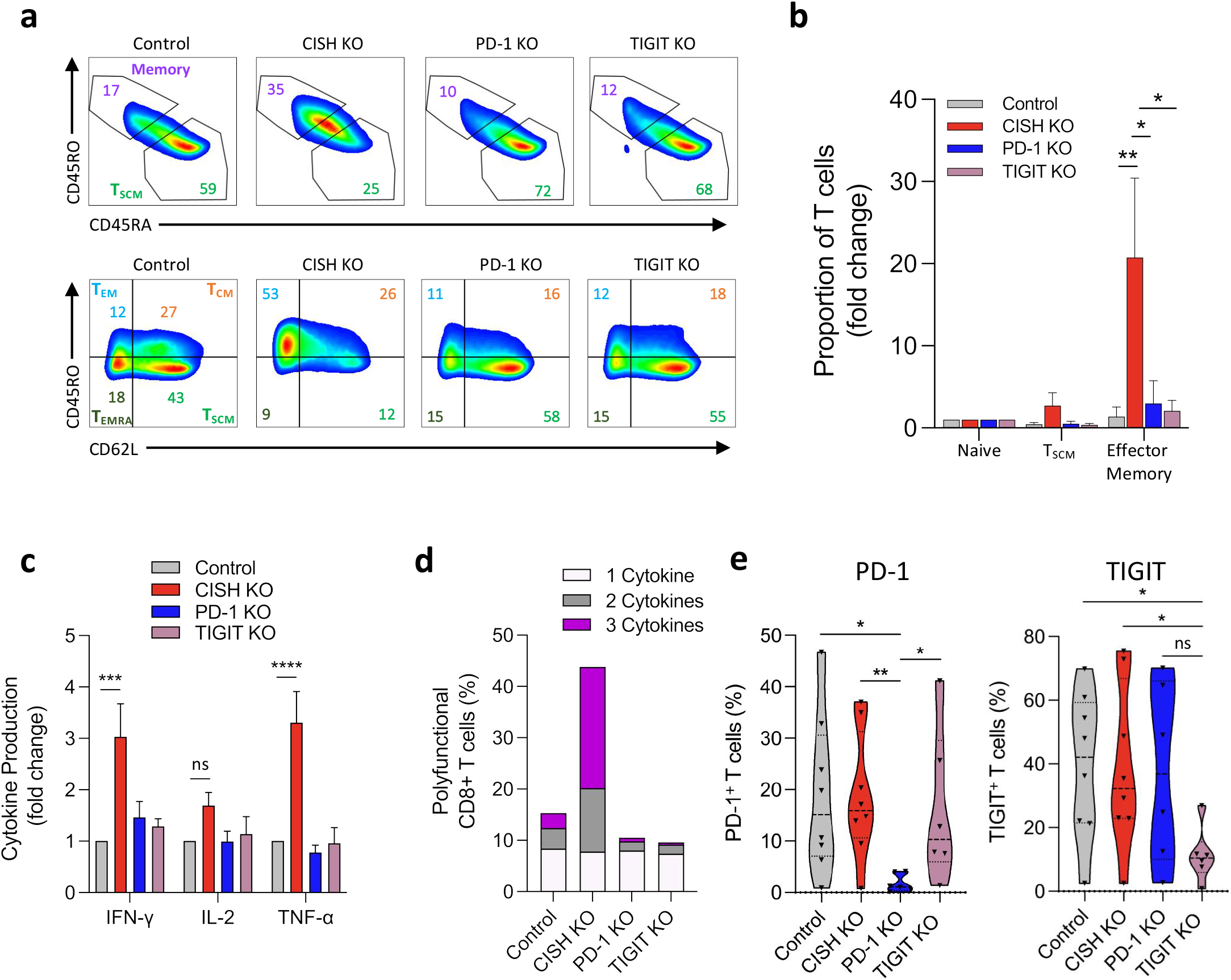
PD-1 or TIGIT disruption does not enhance in T cell effector functions upon TCR signaling. **(a)** The increase in T cell memory formation measured by flow cytometry after anti-CD3/CD8 stimulation, in particular effector memory cells, observed by knockout of CISH is not seen by either PD-1 or TIGIT inactivation. **(b)** Quantification of changes in memory phenotypes in CD8^+^ T cells in control or inactivation of CISH, PD-1, or TIGIT. **(c)** Knockout of CISH significantly increases the magnitude of effector cytokine production measured by ICS after anti-CD3/CD8 stimulation, whereas PD-1 or TIGIT knockout results in cytokine production similar to control T cells. **(d)** Contrary to CISH knockout CD8^+^ T cells, PD-1 or TIGIT knockout does not enhance cytokine polyfunctionality. **(e)** knockout of PD-1 has no impact on expression of TIGIT in anti-CD3/CD8 stimulated CD8^+^ T cells, and similarly knockout of TIGIT does not impact expression of PD-1. Statistical significance was determined by either student t test or ANOVA for repeated measures, *P >0.05, **P>0.01, ***P>0.001, ****P>0.0001. All data are representative of three independent experiments. Error bars represent mean +/− SEM.

### CRISPR inactivation of CISH enhances antigen-specific T cell cytolysis of tumor cells

To resolve the impact of CISH disruption on neoantigen-specific tumor cytolysis, we developed a real-time kinetic cancer cell cytolysis assay using an automated xCELLigence Real-Time Cell Analysis (RTCA) instrument. By co-culturing T cells expressing the KRAS(G12D)-specific TCR with HLA-C*08:02^+^ cells pulsed with either the KRAS(G12D) minimal epitope or the corresponding wild type (WT) sequence, we could reveal an effective level of antigen-specific killing, defining a robust assay window to measure the impact of immune checkpoint gene inactivation on antigen-specific cytolysis (**Fig. 4a**). Killing of antigen-bearing cells by CISH-deficient, KRAS(G12D) TCR expressing CD8^+^ T cells was significantly elevated, both in rapidity and in overall magnitude of response, at timepoints throughout the 5-day co-culture (**Fig. 4b and c**). Co-incubation of CRISPR edited CD8^+^ T cells with the antigen-bearing target cells also lead to a measurable antigen-specific cell lysis at 16 and 48 hours when measured by an orthogonal assay utilizing apoptosis-specific dyes, which was significantly increased by inactivation of CISH (**Fig. 4d**). These cytolytic data demonstrate that inactivation of CISH significantly elevates the neoantigen-specific killing of target cells.

**Figure 4:**
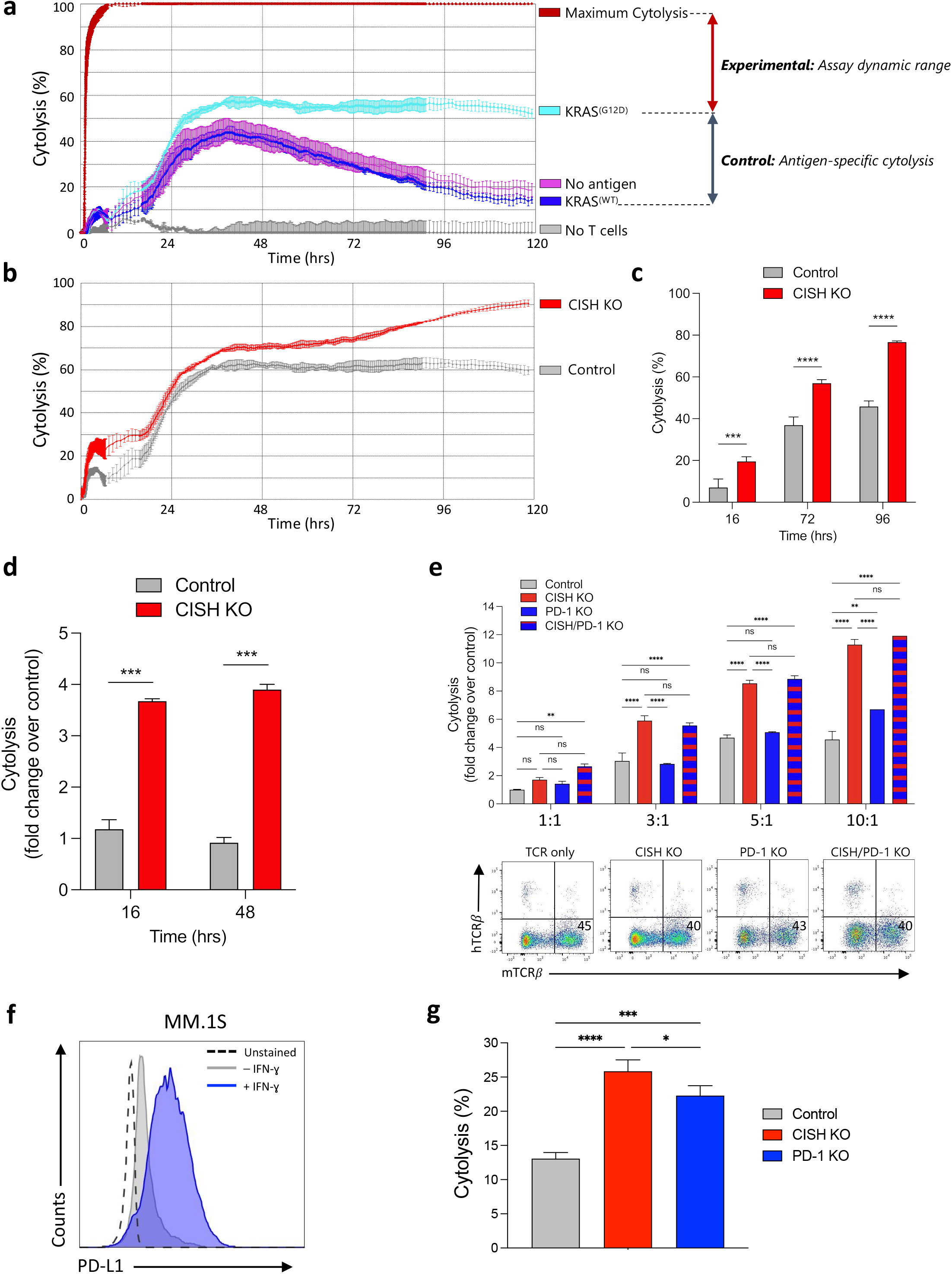
The increased antigen-specific cancer cell killing by CISH disrupted T cells is elevated above that seen by PD-1 deficient T cells. **(a)** A kinetic tumor impedance assay using the xCELLigence system enables real-time measurement of KRAS G12D antigen-specific killing of peptide-pulsed COS-7 cell by CRISPR edited CD8^+^ T cells. **(b)** Increase in the magnitude of antigen-specific cell cytolysis of antigen pulsed COS-7 target cells in the absence of CISH. **(c)** Quantification of 16h, 48h, and 72h timepoints from the cytolysis assay shows a significantly higher cytolytic response is observed at all timepoints for CD8^+^ T cells lacking CISH compared to Control. **(d)** Similar results were observed in an orthogonal assay for cytolysis using Cytox Green as an indicator for cell death when coculturing peptide pulsed COS-7 target cells with control and CISH-edited CD8^+^ T cells, in the presence of cancer-specific KRAS G12D antigen. **(e)** CISH inactivation enhances antigen-specific cytolysis compared to control T cells and loss of PD-1 shows no improvement in cytolysis of COS-7 cells naturally lacking PD-L1. Targeting both CISH and PD-1 shows no synergistic effect, emphasizing the ligand-dependency of PD-1 and ligand independency of CISH in this cellular model. Control condition reflects the KRAS G12D TCR knock-in only. Successful integration of the KRAS G12D TCR in these different gene-edited conditions as well as knockout of the endogenous TCR is confirmed by flow cytometry (panels below). **(f)** The myeloma MM.1S cell line shows a detectible PD-L1 expression which is robustly increased in response to INF-*γ* stimulation. **(g)** When coculturing gene-edited CD8^+^ T cells with MM.1S cancer cells for 16 hours, both CISH knockout and PD-1 knockout T cell enhance the proportion of apoptotic MM.1S cells (measure by Annexin-V staining) compared to control T cells. Statistical significance was determined by either student t test or ANOVA for repeated measures, *P >0.05, **P>0.01, ***P>0.001, ****P>0.0001. All data are representative of at least three independent experiments. Error bars represent mean +/− SEM.

### The enhancement in T Cell function by PD-1 inactivation is only revealed in the presence of PD-L1

We next sought to compare the enhanced cytolysis seen with CISH knockout to T cells lacking PD-1 in the neoantigen-specific killing assay. Surprisingly, we did not observe any change in the killing capacity of PD-1-deficient T cells towards the neoantigen expressing target cells when compared to control cells at any effector to target ratio tested **(Fig. 4e)**. When PD-1 knockout was combined with CISH knockout in the same T cell pool, no additional benefit to PD-1 inactivation was seen above the elevated cytolysis due to CISH-deficiency alone **(Fig. 4e)**. These data suggested that unlike T cells lacking CISH, these conditions of neoantigen-specific TCR stimulation were insufficient for PD-1 disruption to benefit the cytolytic response.

We reasoned that absence of increased cytolysis when PD-1 is knocked out could be due to a requirement for PD-L1 ligand. COS-7 is a monkey kidney fibroblast cell line that does not express high levels of PD-L1; thus, PD-1 knockout T cells were unlikely to have an advantage in eliciting a stronger cytolytic response^29, 30^. To test the requirement for PD-L1, we selected MM.1S, a multiple myeloma cell line that expresses high levels of PD-L1^31, 32^ **(Fig. 4f)**. When CISH knockout CD8^+^ T cells were co-cultured with MM.1S cells for 16 hours, we observed a significantly elevated level of cytolysis compared with control T cells. Similarly, we observed that T cells deficient for PD-1 also exhibited increased cytolysis, albeit at a lower magnitude compared with CISH-deficient T cells **(Fig. 4g)**. These data confirmed that PD-1 editing only leads to enhanced cytolysis when PD-L1 is expressed by a tumor cell line. By contrast, CISH inactivation enhances the cytolytic activity of CD8^+^ T cells regardless of immune checkpoint ligand expression.

### T cell cytolysis of PD-L1 expressing tumor cells is enhanced by PD-1 knockout and further elevated in CISH-deficient T cells

We next analyzed cytolysis of a high PD-L1 expressing cancer cell line by immune checkpoint knockout T cells. Expression of PD-L1 in different tumor lines was analyzed and the ES-2 ovarian clear cell carcinoma line was selected based on its highest expression of PD-L1 **(Fig. 5a)**. This cell line is WT KRAS but expresses the HLA-C*08:02 allele, as confirmed by MHC allele sequencing and from published HLA haplotype data **(Supplementary Fig. S1)**^33, 34, 35, 36^. Thus, in addition to expressing high levels of PD-L1, the ES-2 cell line can be used to present the HLA-C*08:02 restricted KRAS(G12D) peptide to the recombinant TCR expressing T cells when pulsed onto the cell line. To compare the requirement of PD-1 signaling in T cell cytolysis within the same cell type, CRISPR was used to knockout the PD-L1 and PD-1 ligand 2 (PD-L2) genes in the ES-2 line **(Fig. 5b)**. PD-L2 is expressed on antigen-presenting cells and certain tumors, including ovarian cancers. Like PD-L1, PD-L2 has also been shown to bind PD-1 and inhibit TCR-mediated proliferation and cytokine production^37^. We first analyzed the cytolysis of ES-2 cells by control CD8^+^ T cells expressing the KRAS(G12D) TCR and observed antigen-specific killing of cells presenting the KRAS(G12D) neoantigen. **(Fig. 5c)**. CISH-deficient T cells showed a significant increase in antigen-specific cytolysis over control T cells of both the parental ES-2 and the PD-L1/PD-L2 knockout cells **(Fig. 5c & d)**. While cytolysis of the high PD-L1/PD-L2 ES-2 cells by PD-1 knockout T cells showed an increase over control T cells, PD-1 knockout T cells did not elevate the overall killing of PD-L1/PD-L2 deficient ES-2 cells **(Fig. 5e & f)**. Despite enhancing cytolysis of the PD-L1/PD-L2 expressing cancer cells over control T cells, editing of PD-1 was less effective than CISH inactivation.

**Figure 5:**
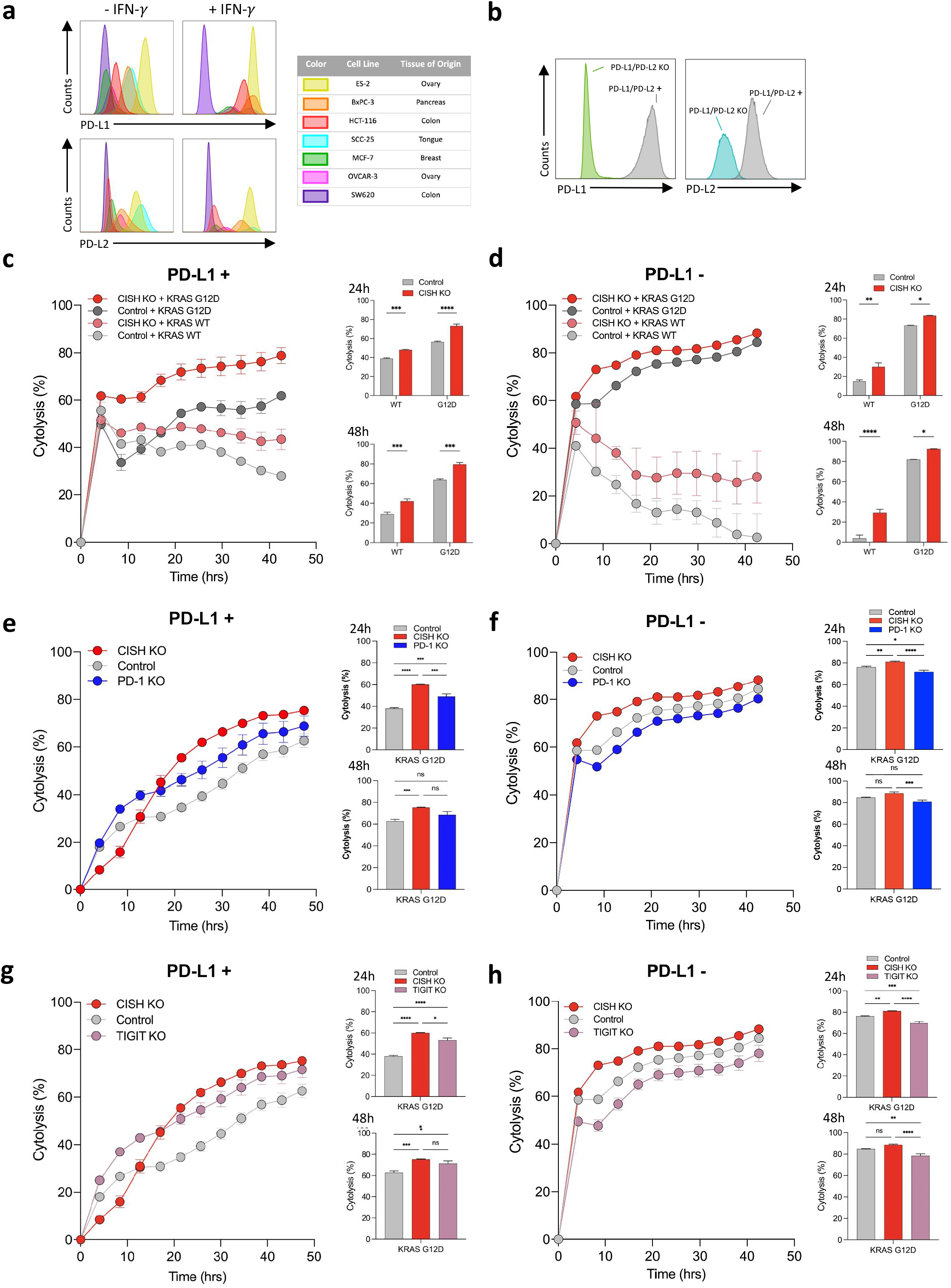
Enhanced antigen-specific cytolysis of PD-1 and TIGIT-deficient T cells is dependent on PD-L1 expression on the cancer cells, whereases elevated cytolysis by CISH inactivation is ligand-independent. **(a)** Human cancer cell lines exhibit varying expression of functional PDL-1 and PDL-2 receptors as shown by upregulation upon treatment with INF-*γ* for 48 hours. Human cancer cell lines evaluated include ES-2 (yellow), BxPC-3 (orange), HCT-116 (red), SCC-25 (blue), MCF-7 (green), OVCAR-3 (pink), and SW620 (purple). **(b)** Sequenced haplotype of HLA-C alleles for each cell line evaluated. **(c)** Loss of expression of PD-L1 and PD-L2 on ES-2 cells engineered by CRISPR. **(d)** CISH disruption enhances antigen-specific T cell cytolysis of KRAS G12D antigen-pulsed ES-2 human cancer cells. **(e)**. The same enhanced cytolysis by CISH inactivation is observed against the PD-L1/PD-L2 knockout ES-2 cells. **(f)** PD-1 knockout results in an increase in antigen-specific ES-2 cell cytolysis, **(g)** but no significant overall increase in cytolysis towards the PD-L1/PD-L2 knockout ES-2 cells, indicating a requirement for the ligands to be present to reveal a cytolytic benefit for PD-1 inactivation. The elevated cytolysis of ES-2 cells by PD-1 KO T cells is lower than observed with CISH KO T cells. **(h-i)** Similar results are observed with TIGIT deficiency in CD8^+^ T cells improving cytolysis in the setting of PD-L1/PD-L2 expression but showing no benefit when these ligands are absent on the ES-2 cells. Antigen-specific cytolysis is elevated by T cells lacking CISH over TIGIT regardless of PD-L1/PD-L2 expression on the cancer cells. Statistical significance was determined by either student t test or ANOVA for repeated measures, *P >0.05, **P>0.01, ***P>0.001, ****P>0.0001. All data are representative of at least three independent experiments. Error bars represent mean +/− SEM.

Analysis of ES-2 cell cytolysis by TIGIT knockout T cells revealed a similar pattern to that seen by inactivation of PD-1, with TIGIT-deficient T cells enhancing cytolysis of ES-2 parental cells significantly over control T cells and a lack of improvement in killing of ES-2 cells lacking PD-L1/PD-L2 **(Fig. 5g & h)**. The dependency of PD-L1/PD-L2 expression on TIGIT regulation of cytolysis may not be surprising, as recent evidence suggests T cell functional and anti-tumor responses regulated by TIGIT and PD-1 appear to be overlapping and co-dependent^38, 39^. Despite significantly elevated cytolysis by TIGIT knockout T cells over control, CISH-deficient T cells lead to the highest increase in antigen-specific cytolysis of these ovarian cancer cells **(Fig. 5g & h)**. Collectively, these data indicate an essential requirement for PD-L1 expression by cancer cells for any demonstrable effect on anti-tumor responses by PD-1 knockout and TIGIT knockout T cells. By contrast, CISH inactivation showed superior improvements in cytolysis irrespective of PD-L1/PD-L2 expression.

### Enhanced anti-tumor cytolysis by CISH-deficient T cells is synergistic in combination to PD-1 or TIGIT immune checkpoint knockout

Finally, given the ligand restriction of PD-1 signaling and expected distinct and non-redundant role with CISH-mediated abrogation of proximal TCR signaling, we evaluated the potential synergy of PD-1 and CISH knockout in the context of neoantigen-specific cancer cell killing. To this end, we performed multiplex CRISPR engineering in KRAS(G12D) TCR targeted CD8^+^ T cells to knockout CISH in combination with either PD-1 or TIGIT in the same pool of T cells and analyzed cytolysis of the ES-2 parental and ES-2 PD-L1/PD-L2 knockout cancer cells. Combined inactivation of CISH and PD-1 resulted in a significantly elevated killing of the parental ES-2 cell lines above either immune checkpoint alone at all timepoints measured **(Fig. 6a)**. This combined efficacy was only evident when PD-L1/2 ligands were present, as the CISH plus PD-1 knockout T cells showed no further increase in killing of ES-2 cells lacking PD-L1/PD-L2 above CISH knockout alone.

**Figure 6:**
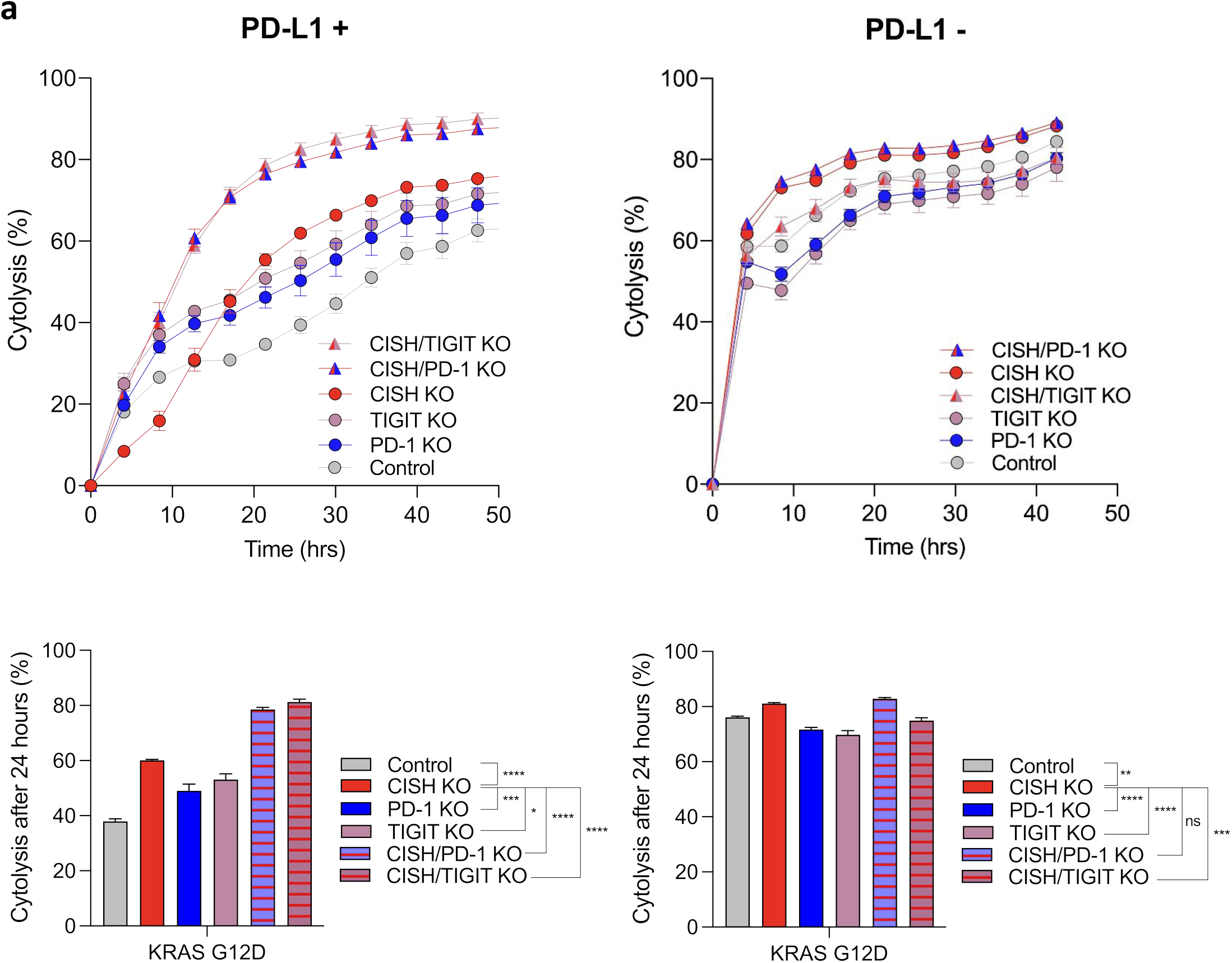
Inactivation of CISH and PD-1/TIGIT synergize to maximize the increase in antigen-specific cancer cell cytolysis. **(a)** Disruption of CISH in combination with PD-1 or TIGIT knockout both show maximum and synergistic levels of antigen-specific cytolysis towards the high PD-L1/PD-L2 ES-2 myeloma cells. This synergy is only observed toward cancer cells harboring functional PD-L1 and PD-L2 and not CRISPR engineered cells lacking these two ligands, emphasizing the ligand-dependency of PD-1 (and TIGIT) and superior, ligand independency of CISH in this cellular model. Statistical significance was determined by either student t test or ANOVA for repeated measures, *P >0.05, **P>0.01, ***P>0.001, ****P>0.0001. All data are representative of at least three independent experiments. Error bars represent mean +/− SEM.

Like PD-1, knockout of TIGIT in combination with CISH inactivation also showed an enhanced level of neoantigen-specific cytolysis of the PD-L1/PD-L2 expressing cancer cells **(Fig. 6a)**. Again, improved killing was only evident in the presence of PD-L1/PD-L2, further demonstrating that cell surface immune checkpoints such as PD-L1 and TIGIT can only add additional cytolytic efficacy to CISH-deficient T cells in a context where tumor cells express their ligands.

## DISCUSSION

Tumor resident antigen-specific T cells, such as neoantigen reactive TIL, can recognize and clear cancer cells, although the clinical efficacy remains promising yet inconsistent with or without combined immune checkpoint inhibition^40, 41^. The TME exerts complex and mostly understudied mechanisms for suppressing T cell function and the upregulation of cell surface immune checkpoint proteins, reduced MHC expression on cancer cells, and low antigen density are only some of the main extrinsic factors contributing to a weakened and short-lived cytolytic T cell response after TCR activation^42, 43, 44^.

MAb-based therapies for inactivating classical cell-surface immune checkpoints such as PD-1/PD-L1, CTLA-4, and possibly TIGIT can help to overcome some of the suppressive effects of the TME on cancer neoantigen-specific T cells and have shown promising clinical outcomes in a subset of patients^45^. However, the requirement for the cancer to express high levels of PD-L1 for mAb blockade to have any meaningful clinical efficacy, and the heterogeneity in PD-L1 expression between cancer types and individuals, restricts the efficacy of this therapeutic approach to a relatively restricted subset of responsive patients^46^.

A new class of intra-cellular immune checkpoints, exemplified by CISH, have the potential to overcome this limit of ligand-dependency and have the potential to enhance the anti-tumor functions of T cells against any cancer in a PD-L1 agnostic manner. Recent studies have highlighted that CISH is highly expressed in activated T cells and TILs isolated from patient tumors and demonstrate the important role CISH plays in negatively regulating TCR avidity, tumor cytolysis and neoantigen recognition^20, 22, 23^. Furthermore, the inactivation of CISH in human TIL resulted in improved antigen-specific activation and unmasked reactivity against shared neoantigens, suggesting that ablation of CISH within the TME may help cancer-specific T cells to overcome T cell intrinsic suppression of the cytolytic response and augment the anti-cancer activity of these reactive cells. The additional finding of increased PD-1 expression in CISH-deficient T cells, and a synergistic response of combined CISH and PD-1 inactivation in a murine melanoma model, warranted further investigation of the comparison and combination of these non-redundant immune checkpoint pathways^20^.

In the current study we build upon recent findings that demonstrate the role of CISH in modulating T cell anti-tumor functions, neoantigen reactivity, and cytolytic effector programs by evaluating the impact of CISH inactivation in antigen-specific, anti-tumor T cell functions in comparison and combination to PD-1 and TIGIT. We developed an optimized CRISPR/Cas9 editing strategy that enables efficient simultaneous genetic disruption of multiple immune checkpoint genes in human T cells while concurrently targeting the endogenous TCR locus to stably integrate and express a recombinant TCR specific to the human shared neoantigen, KRAS(G12D).

Our findings demonstrate that CISH knockout results in a significant enhancement in TCR stimulated T_EM_ cell formation and cytokine production, highlighting the important role this target plays downstream of the activated TCR. Surprisingly, our experiments did not show a similar significant enhancement of these functional T cell responses when either PD-1 or TIGIT was inactivated by CRISPR. This finding suggests that unlike CISH, TCR stimulation alone may not be sufficient to reveal the benefits of disruption of PD-1 and TIGIT signaling pathways in T cells.

To accurately compare the impact of genetic disruption of immune checkpoint genes on anti-tumor activity *in vitro*, we developed an evaluation platform where CRISPR-edited T cells can be tested for their capacity to kill neoantigen-bearing tumor cells in a sensitive and real-time assay. These tumor cell killing assays demonstrated that cytolysis of antigen-expressing tumor cells by CISH-deficient T cells was significantly elevated over control T cells. Furthermore, the finding that CISH knockout led to elevated tumor cell killing in all conditions and biological donors tested, regardless of the cell type or PD-L1expression, supports the notion that CISH, by virtue of being intra-cellular and a key regulator of proximal TCR signaling, operates to control T cell responses in a ligand-independent manner. Antigen-specific TCR stimulation alone was not sufficient for PD-1 inactivation to benefit anti-tumor responses. Further, we demonstrated that expression of PD-L1/L2 is required for PD-1 knockout T cells to enhance cytolysis.

Interest in TIGIT as an anti-cancer target has increased recently and anti-TIGIT mAbs are now being evaluated in early clinical trials with modest yet evolving data^47, 48^. While more is known regarding the biology of PD-1 and its ligand interactions, ligands of TIGIT have been identified as poliovirus receptor (PVR), Nectin2, Nectin3 and Nectin4 and have been shown to be expressed by tumor cells and antigen-presenting cells within the TME^49, 50, 51, 52^. Our data showing TIGIT-deficient T cells induced a similar cytolytic response to PD-1 knockout T cells, whereby an enhanced level of tumor cell killing was revealed in the absence of PD-L1 signaling, suggests a potential interdependency on PD-1-mediated inhibition of T cell activation and function.

The precision and efficiency of multiplex CRISPR editing enables the inactivation of multiple genes within the same T cell and enables us to evaluate the combined genetic disruption of both CISH and PD-1 or TIGIT. The enhancement of neoantigen-specific tumor cell killing that we observed suggests that CISH and PD-1 independently regulate T cell function using non-redundant signaling pathways. These findings highlight the promising notion of combination immune checkpoint inhibition for enhancing anti-cancer response that may leverage the distinct pathways of both intra-cellular and cell surface immune checkpoint targets. As predicted, this additive response was only seen against tumor cells expressing high levels of PD-L1, whereby PD-1 knockout or TIGIT knockout appears to bypass the suppressive effects of PD-L1 expression and enhance cytolysis above and beyond the increased killing observed with CISH-inactivation alone.

The ideal attributes of immune checkpoint targets for efficacy in solid cancers can be considered in terms of the effectiveness of tumor cell killing and accessibility for precise drugging and thus therapeutic inhibition. As summarized in **Table 1**, these attributes show distinct differences between cell surface immune checkpoints PD-1 and TIGIT, and the intra-cellular immune checkpoint CISH. While disruption of all these targets can improve neoantigen-specific tumor cell lysis as demonstrated in this study, the ligand independency of CISH offers the potential for broadening immune checkpoint therapies against any solid cancer. The durable clinical benefit of anti-PD-1 immunotherapy is now well established and recent data suggests anti-tumor efficacy is also achievable through TIGIT blockade in combination to anti-PD-L1 therapy^47^. The intra-cellular nature of CISH makes conventional immune checkpoint inhibition using a mAb-based therapy challenging. While direct intra-cellular protein drugging modalities for CISH may one day be possible, precision genetic engineering has now enabled the efficacy of the CISH immune checkpoint to be objectively evaluated in ongoing clinical trials.

**Table 1:**
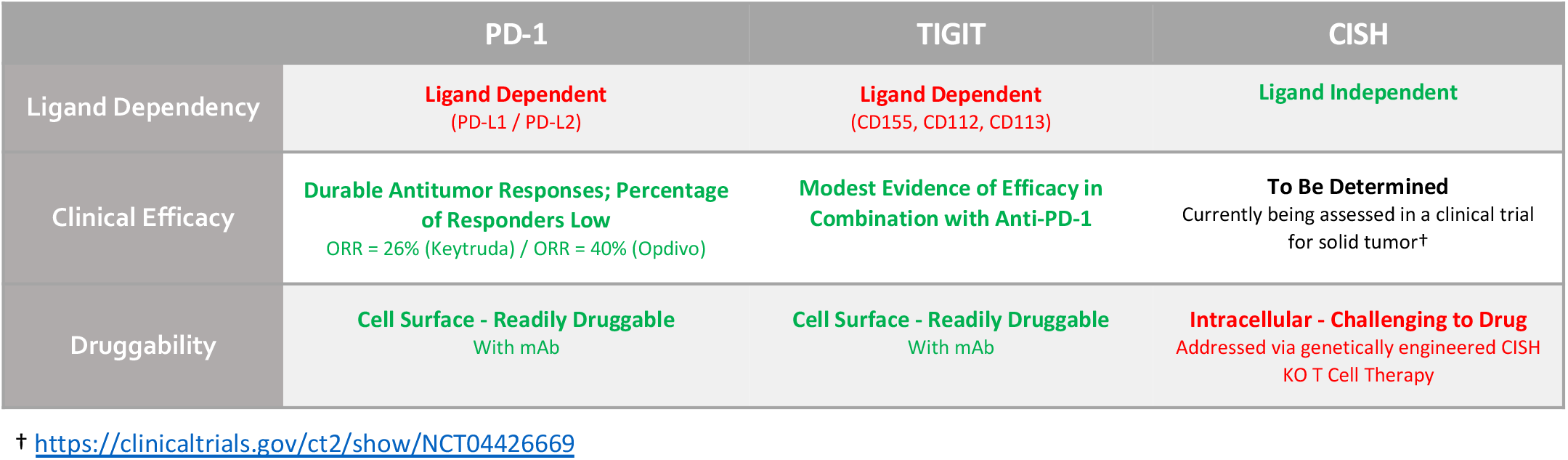
The attributes of intra-cellular CISH inhibition in human T cells in comparison to cell surface immune checkpoints PD-1 and TIGIT.

|  | PD-1 | TIGIT | CISH |
| --- | --- | --- | --- |
| Ligand Dependency | Ligand Dependent<br>(PD-L1 / PD-L2) | Ligand Dependent<br>(CD155, CD112, CD113) | Ligand Independent |
| Clinical Efficacy | Durable Antitumor Responses; Percentage of Responders Low<br>ORR = 26% (Keytruda) / ORR = 40% (Opdivo) | Modest Evidence of Efficacy in Combination with Anti-PD-1 | To Be Determined<br>Currently being assessed in a clinical trial for solid tumors† |
| Druggability | Cell Surface - Readily Druggable<br>With mAb | Cell Surface - Readily Druggable<br>With mAb | Intracellular - Challenging to Drug<br>Addressed via genetically engineered CISH KO T Cell Therapy |

† https://clinicaltrials.gov/ct2/show/NCT04426669

The finding that concurrent inactivation of CISH and PD-1 can act together to further improve the tumor-specific cytolytic potential of T cells offers a compelling prospect for a dual-therapeutic approach to target both immune checkpoint genes in a hope to engender a T cell therapy with a durable and complete anti-cancer response. The enhanced anti-tumor response observed with CISH-deficient T cells in this and other published reports positions CISH as a next-generation intra-cellular immune checkpoint target that may have meaningful clinically efficacy in the setting of a broad cross-section of solid cancers, irrespective of the presence of PD-L1/PD-L2 or other immunosuppressive ligands.

## METHODS

### PBMC samples and isolation of CD8^+^ T cells

Peripheral blood mononuclear cells were obtained from anonymized healthy individuals (Caltag Medsystems, Tissue Solutions Ltd and Precision for Medicine, Inc.) and obtained, handled, and stored in accordance with the Human Tissue Authority UK regulations. Total CD8^+^ T cells were isolated from unfractionated PBMCs using the EasySep Human CD8^+^ T Cell Isolation Kit (Stem Cell Technologies) with a DynaMag-2 magnet (ThermoFisher Scientific) according to the manufacturer’s guidelines. The ratio of CD8:CD4 and viability of isolated T cells was assessed regularly using flow cytometry.

### Expansion of CD8^+^ T cells

Isolated human CD8^+^ T cells were cultured in X-VIVO-15 Basal Media (Lonza) supplemented with 10% Human AB Serum Heat Inactivated (Sigma), 300IU/ml Recombinant Human IL-2, 5ng/ml Recombinant Human IL-7, and 5ng/ml Recombinant Human IL-15 (all Peprotech) and 10mM N-Acetyl-L-cysteine (Sigma) and cultured in a 37°C, 5% CO_2_ cell culture incubator. Media was replaced every 2-3 days with fresh complete media including cytokines.

### Cell Lines and Culturing

All cell lines used for this study were purchased from ATCC and cultured in their recommended media formulations and growth conditions. Cells were kept at sub-confluent densities and regularly tested for mycoplasma. SV40-transformed COS-7 cells were engineered to express a human HLA C*08:02 allele to enable presentation of pulsed KRAS wildtype and mutant peptides. HLA-A/B/C allele typing for each cell line was performed by MC Diagnostics Ltd (UK).

### sgRNA Design

sgRNAs targeting *TRAC, CISH, PDCD1, VSIG9, PDCD1LG1 (PD-L1)*, and *PDCD1LG2* (*PD-L2*) were designed using various online resources. Up to 6 sgRNAs per target gene were tested and the most efficient sgRNA was selected containing 2′-O-methyl and 3′ phosphorothioate modifications to the first three 5′ and the last three 3′ nucleotides (Synthego).

### Production of AAV-mediated TCR-Knock-in and checkpoint-knockout CD8^+^ T cells using CRISPR/Cas9

CD8+T cells were stimulated using anti-CD3/CD28 Dynabeads (Life Technologies) in complete T cell media and under normal growth conditions for 48-72 hours prior to electroporation. T cells were electroporated in Neon Buffer T with 15*μ*g Cas9 mRNA (TriLink) and 10*μ*g total sgRNA (Synthego) using the Neon electroporator (3×10^6^ cells per 100*μ*l tip) (Life Technologies) using parameters 1400V, 10ms, 3 pulses. To achieve targeted recombinant TCR integration into the *TRAC* locus, rAAV6 was added to CD8^+^ T cells 3-5 hours after electroporation at an MOI of 1×10^6^ particles per cell. Viral rAAV6 particles were produced by Vigene Biosciences or PackGene. Electroporated T cells were recovered in complete T cell media at a density of 1×10^6^ cells per ml and allowed to rest for 48 hours before subsequent analysis.

### Analysis of Gene Knockout Efficiency on DNA Level

Primers for PCR were designed to amplify a 600-900 base pair region surrounding the sgRNA target site. A minimum of 24 hours after electroporation, genomic DNA was extracted from CD8^+^ T cells using the DirectPCR Lysis solution (Viagen Biotech) containing Proteinase K and target regions were amplified by PCR using the GoTaq G2 PCR mastermix (Promega). Correct and unique amplification of the target regions was verified by agarose gel electrophoresis before purifying PCR products using the QIAquick PCR Purification Kit (Qiagen). For analysis by TIDE, PCR amplicons were Sanger sequenced (Eurofins or Genewiz), and paired .ab1 files of control versus edited samples were analyzed using Synthego’s ICE tool (https://ice.synthego.com).

### Immunoblot analysis

*Western* blot analysis was performed using standard protocols. In brief, cells were harvested and washed once in ice-cold PBS and then lysed in 1X RIPA Buffer containing 1X Protease Inhibitors on ice for 10 minutes. Cells were then centrifuged in a table-top centrifuge at 14,000 rcf for 20 min at 4°C to pellet cell debris. Proteins were separated on a 4–12% SDS-PAGE gel followed by standard immunoblot analysis using anti–CISH (Cell Signaling, Clone D4C10, 1:2000) and Vinculin (Cell Signaling, Clone EPR8185, 1:5000). Detection of proteins was performed using secondary antibodies conjugated to horseradish peroxidase-HRP and the SuperSignal West Pico Plus chemiluminescent substrate (Thermo Scientific-Pierce).

### Flow cytometry analysis of T cell phenotypes

For flow cytometric analysis of the CRISPR edited T cell phenotypes and cell surface marker expression, cells were harvested from culture plates and washed using FACS Buffer containing PBS with 0.5% Bovine Serum Albumin (Thermo Scientific) and were then stained with monoclonal antibodies specific for CD8 (HIT8A, 1:100), CD4 (OKT4, 1:100), HLA-DR (L243 1:80), LAG-3 (11C3C65, 1:80), TIGIT (VSTM3, 1:40), CD45RO (UCHL1, 1:40), CD45RA (HI100, 1:80), TIM3 (F38-2E2, 1:40), CD62L (DREG-56, 1:40), CD57 (QA17A04, 1:80), PD-1 (EH12.1, 1:40), OX-40 (Ber-ACT35, 1:40), CD25 (MA2-51, 1:40), 41BB (4B4-1, 1:40), (Biolegend) or specific for CD8 (RPA-T8, 1:100) (BD Bioscience) and CD3 (UCHT1, 1:100) (ThermoFisher). Live/Dead Fixable Dead Cell Stains (Invitrogen) or SYTOX Blue Dead Cell Stain (Invitrogen) were included in all experiments to exclude dead cells. All samples were acquired on a Fortessa flow cytometer (BD Bioscience), and data was analyzed using FlowJo 10 software (BD Biosciences).

### Intra-cellular cytokine staining

Cells were stimulated for a total of 6 hours with human T-activator anti-CD3/CD28 Dynabeads (ThermoFisher) stimulation with GolgiStop solution being added for a total of 5 hours block intra-cellular protein transport (BD Bioscience). As a positive control for cytokine production, a pool of T cells was stimulated for 6 hours with 50ng/ml PMA and 1*μ*g/ml Ionomycin (Sigma). T cells were then harvested and washed with FACS Buffer and stained for surface markers followed by fixation and permeabilization using BD Cytofix/Cytoperm Fixation/Permeabilization Solution (ThermoFisher) before proceeding with intra-cellular cytokine staining using antibodies specific for INF-*γ* (4S.B3, 1:40) (Biolegend) IL-2 (MQ1-17H12, 1:40) (BD Bioscience), or TNF-*α* (MAb11, 1:40) (ThermoFisher). All samples were acquired on a Fortessa flow cytometer (BD Bioscience), and data was analyzed using FlowJo 10 software (BD Biosciences).

### Realtime Cytolysis Assay (RTCA)

Cytolysis assays were carried out with the xCELLigence RTCA SP platform (Acea Bioscience/Agilent) based on electrical impedance resulting in a cell index (CI) value. Background measurements were taken with media only before seeding cells. Adherent COS-7 or ES-2 tumor cells were then plated in a 96-well RTCA View plate at a pre-determined density per well to reach a linear growth time phase after roughly 14-18 hours of culture and incubated overnight at 37°C and 5% CO2 in their respective complete growth medium. The next day, cancer cells were pulsed with mutant (G12D) or wildtype (WT) peptides for 2 hours and then washed prior to the addition of different knockout T cells or Control T cells. T cells were added at indicated effector to target cell ratios (E:T) and containing the respective gene edits. Cytolysis assays were run for up to 90 hours undisturbed with measurements taken every 2-10 minutes. Data was analyzed using RTCA software and plotted as % Cytolysis calculated as (impedance of target cells without effector cells – impedance of target cells with effector cells) x100 divided by impedance of target cells without effector cells. Controls include background measurements as well as a negative control containing target cells only as well as a positive control containing target cells receiving 2.5% Triton-x solution for maximum cytolysis.

### Statistical analyses

Statistical differences between two sample groups, where appropriate, were analyzed by a standard Student’s two-tailed, non-paired, t-test and between three or more sample groups using two-way or three-way ANOVA using GraphPad Prism Software version 9. P values are included in the figures where statistical analyses have been carried out.

## Ethics declarations

The authors declare no competing interests.

## ACKNOWLEDGEMENTS

This study was supported by Intima Bioscience, NIH R37 CA259177 (C.A.K.), and NIH P30 CA008748 (C.A.K.).

## FIGURES

**Supplementary Figure S1: Sequenced haplotype of HLA-C alleles for each cancer cell line evaluated**.

## REFERENCES

1. Heslop, H.E. et al. Long-term restoration of immunity against Epstein-Barr virus infection by adoptive transfer of gene-modified virus-specific T lymphocytes. Nat Med 2, 551–555 (1996).

2. Campillo-Davo, D., Flumens, D. & Lion, E. The Quest for the Best: How TCR Affinity, Avidity, and Functional Avidity Affect TCR-Engineered T-Cell Antitumor Responses. Cells 9 (2020).

3. Chandran, S.S. & Klebanoff, C.A. T cell receptor-based cancer immunotherapy: Emerging efficacy and pathways of resistance. Immunol Rev 290, 127–147 (2019).

4. Darvin, P., Toor, S.M., Sasidharan Nair, V. & Elkord, E. Immune checkpoint inhibitors: recent progress and potential biomarkers. Exp Mol Med 50, 1–11 (2018).

5. Song, M., Chen, X., Wang, L. & Zhang, Y. Future of anti-PD-1/PD-L1 applications: Combinations with other therapeutic regimens. Chin J Cancer Res 30, 157–172 (2018).

6. Munhoz, R.R. & Postow, M.A. Clinical Development of PD-1 in Advanced Melanoma. Cancer J 24, 7–14 (2018).

7. Rotte, A. Combination of CTLA-4 and PD-1 blockers for treatment of cancer. J Exp Clin Cancer Res 38, 255 (2019).

8. Sun, L. et al. Clinical efficacy and safety of anti-PD-1/PD-L1 inhibitors for the treatment of advanced or metastatic cancer: a systematic review and meta-analysis. Sci Rep 10, 2083 (2020).

9. Upadhaya, S. et al. Combinations take centre stage in PD-1/PDL1 inhibitor clinical trials. Nat Rev Drug Discov (2020).

10. Seidel, J.A., Otsuka, A. & Kabashima, K. Anti-PD-1 and Anti-CTLA-4 Therapies in Cancer: Mechanisms of Action, Efficacy, and Limitations. Front Oncol 8, 86 (2018).

11. Riley, J.L. PD-1 signaling in primary T cells. Immunol Rev 229, 114–125 (2009).

12. He, X. & Xu, C. Immune checkpoint signaling and cancer immunotherapy. Cell Res 30, 660–669 (2020).

13. Martin-Ruiz, A. et al. Effects of anti-PD-1 immunotherapy on tumor regression: insights from a patient-derived xenograft model. Sci Rep 10, 7078 (2020).

14. Lee, J. et al. Outstanding clinical efficacy of PD-1/PD-L1 inhibitors for pulmonary pleomorphic carcinoma. Eur J Cancer 132, 150–158 (2020).

15. Nowicki, T.S., Hu-Lieskovan, S. & Ribas, A. Mechanisms of Resistance to PD-1 and PD-L1 Blockade. Cancer J 24, 47–53 (2018).

16. Jenkins, R.W., Barbie, D.A. & Flaherty, K.T. Mechanisms of resistance to immune checkpoint inhibitors. Br J Cancer 118, 9–16 (2018).

17. Sun, C., Mezzadra, R. & Schumacher, T.N. Regulation and Function of the PD-L1 Checkpoint. Immunity 48, 434–452 (2018).

18. Pawelczyk, K. et al. Role of PD-L1 Expression in Non-Small Cell Lung Cancer and Their Prognostic Significance according to Clinicopathological Factors and Diagnostic Markers. Int J Mol Sci 20 (2019).

19. Davis, A.A. & Patel, V.G. The role of PD-L1 expression as a predictive biomarker: an analysis of all US Food and Drug Administration (FDA) approvals of immune checkpoint inhibitors. J Immunother Cancer 7, 278 (2019).

20. Douglas C. Palmer, et al. Internal checkpoint regulates T cell neoantigen reactivity and susceptibility to PD-1 blockade. bioRxiv In press (2020).

21. Yoshimura, A. CIS: the late-blooming eldest son. Nat Immunol 14, 692–694 (2013).

22. Palmer, D.C. et al. Cish actively silences TCR signaling in CD8+ T cells to maintain tumor tolerance. The Journal of experimental medicine 212, 2095–2113 (2015).

23. Guittard, G. et al. The Cish SH2 domain is essential for PLC-gamma1 regulation in TCR stimulated CD8(+) T cells. Sci Rep 8, 5336 (2018).

24. Delconte, R.B. et al. CIS is a potent checkpoint in NK cell-mediated tumor immunity. Nat Immunol 17, 816–824 (2016).

25. Zhu, H. et al. Metabolic Reprograming via Deletion of CISH in Human iPSC-Derived NK Cells Promotes In Vivo Persistence and Enhances Anti-tumor Activity. Cell Stem Cell 27, 224–237 e226 (2020).

26. Miah, M.A. et al. CISH is induced during DC development and regulates DC-mediated CTL activation. Eur J Immunol 42, 58–68 (2012).

27. ClinicalTrials.gov [Internet]. Bethesda (MD): National Library of Medicine (US). 2000 Feb 29 -. Identifier NCT04426669, A Study of Metastatic Gastrointestinal Cancers Treated With Tumor Infiltrating Lymphocytes in Which the Gene Encoding the Intracellular Immune Checkpoint CISH Is Inhibited Using CRISPR Genetic Engineering; 2020 June 11. Available from: https://clinicaltrials.gov/ct2/show/NCT04426669.

28. Tran, E. et al. Immunogenicity of somatic mutations in human gastrointestinal cancers. Science 350, 1387–1390 (2015).

29. Ikebuchi, R. et al. Influence of PD-L1 cross-linking on cell death in PD-L1-expressing cell lines and bovine lymphocytes. Immunology 142, 551–561 (2014).

30. Chen, S. et al. Mechanisms regulating PD-L1 expression on tumor and immune cells. J Immunother Cancer 7, 305 (2019).

31. Zhao, Z. et al. CRISPR knock out of programmed cell death protein 1 enhances anti-tumor activity of cytotoxic T lymphocytes. Oncotarget 9, 5208–5215 (2018).

32. Lanzel, E.A. et al. Predicting PD-L1 expression on human cancer cells using next-generation sequencing information in computational simulation models. Cancer Immunol Immunother 65, 1511–1522 (2016).

33. Boegel, S., Lower, M., Bukur, T., Sahin, U. & Castle, J.C. A catalog of HLA type, HLA expression, and neo-epitope candidates in human cancer cell lines. Oncoimmunology 3, e954893 (2014).

34. Bairoch, A. The Cellosaurus, a Cell-Line Knowledge Resource. J Biomol Tech 29, 25–38 (2018).

35. Nakayama, N. et al. KRAS or BRAF mutation status is a useful predictor of sensitivity to MEK inhibition in ovarian cancer. Br J Cancer 99, 2020–2028 (2008).

36. Tate, J.G. et al. COSMIC: the Catalogue Of Somatic Mutations In Cancer. Nucleic Acids Res 47, D941–D947 (2019).

37. Latchman, Y. et al. PD-L2 is a second ligand for PD-1 and inhibits T cell activation. Nat Immunol 2, 261–268 (2001).

38. Zhang, Q. et al. Blockade of the checkpoint receptor TIGIT prevents NK cell exhaustion and elicits potent anti-tumor immunity. Nat Immunol 19, 723–732 (2018).

39. Johnston, R.J. et al. The immunoreceptor TIGIT regulates antitumor and antiviral CD8(+) T cell effector function. Cancer Cell 26, 923–937 (2014).

40. Spranger, S. et al. Density of immunogenic antigens does not explain the presence or absence of the T-cell-inflamed tumor microenvironment in melanoma. Proceedings of the National Academy of Sciences of the United States of America 113, E7759–E7768 (2016).

41. Creelan, B.C. et al. Tumor-infiltrating lymphocyte treatment for anti-PD-1-resistant metastatic lung cancer: a phase 1 trial. Nat Med 27, 1410–1418 (2021).

42. Xia, A., Zhang, Y., Xu, J., Yin, T. & Lu, X.J. T Cell Dysfunction in Cancer Immunity and Immunotherapy. Front Immunol 10, 1719 (2019).

43. Zhao, Y., Shao, Q. & Peng, G. Exhaustion and senescence: two crucial dysfunctional states of T cells in the tumor microenvironment. Cell Mol Immunol 17, 27–35 (2020).

44. Anderson, K.G., Stromnes, I.M. & Greenberg, P.D. Obstacles Posed by the Tumor Microenvironment to T cell Activity: A Case for Synergistic Therapies. Cancer Cell 31, 311–325 (2017).

45. Egen, J.G., Ouyang, W. & Wu, L.C. Human Anti-tumor Immunity: Insights from Immunotherapy Clinical Trials. Immunity 52, 36–54 (2020).

46. Reck, M. et al. Pembrolizumab versus Chemotherapy for PD-L1-Positive Non-Small-Cell Lung Cancer. N Engl J Med 375, 1823–1833 (2016).

47. Chauvin, J.M. & Zarour, H.M. TIGIT in cancer immunotherapy. J Immunother Cancer 8 (2020).

48. Solomon, B.L. & Garrido-Laguna, I. TIGIT: a novel immunotherapy target moving from bench to bedside. Cancer Immunol Immunother 67, 1659–1667 (2018).

49. Casado, J.G. et al. Expression of adhesion molecules and ligands for activating and costimulatory receptors involved in cell-mediated cytotoxicity in a large panel of human melanoma cell lines. Cancer Immunol Immunother 58, 1517–1526 (2009).

50. Levin, S.D. et al. Vstm3 is a member of the CD28 family and an important modulator of T-cell function. Eur J Immunol 41, 902–915 (2011).

51. Stanietsky, N. et al. The interaction of TIGIT with PVR and PVRL2 inhibits human NK cell cytotoxicity. Proc Natl Acad Sci U S A 106, 17858–17863 (2009).

52. Yu, X. et al. The surface protein TIGIT suppresses T cell activation by promoting the generation of mature immunoregulatory dendritic cells. Nat Immunol 10, 48–57 (2009).

