## Supplementary figure S1 for "Genetic editing of *CISH* enhances T cell effector programs independently of immune checkpoint cell surface ligand expression"

**Supplementary Figure S1:** Sequenced haplotype of HLA-C alleles for each cancer cell line evaluated.

| Cell Line | Cancer Type | HLA-C serotype |
|---|---|---|
| **ES-2** | **Ovarian clear cell adenocarcinoma** | **C*07:01, 08:02** |
| SW1990 | Pancreatic adenocarcinoma | C*01:02, 12:03 |
| BxPC-3 | Pancreatic ductal adenocarcinoma | C*06:02 |
| SCC25 | Tongue squamous cell carcinoma | C*01:01, 07:01 |
| HCT-116 | Colorectal Carcinoma | C*05:01, 07:01 |
